# Targeted Protein O-GlcNAcylation Using Bifunctional Small Molecules

**DOI:** 10.1101/2023.06.09.544275

**Authors:** Bowen Ma, Khadija Shahed Khan, Tongyang Xu, Josefina Xeque Amada, Zhihao Guo, Yu Yan, Alfred Sze-Lok Cheng, Billy Wai-Lung Ng

## Abstract

Protein O-linked β-*N*-acetylglucosamine modification (O-GlcNAcylation) plays a crucial role in regulating essential cellular processes. The disruption of O-GlcNAcylation homeostasis has been linked to various human diseases, including cancer, diabetes, and neurodegeneration. However, there are limited chemical tools for protein- and site-specific O-GlcNAc modification, rendering the precise study of O-GlcNAcylation challenging. To address this, we have developed first-in-class heterobifunctional small molecules, named O-GlcNAcylation targeting chimeras (OGTACs), which enable protein-specific O-GlcNAcylation in cells. OGTACs promote O-GlcNAcylation of proteins such as BRD4, CK2α, and EZH2 *in cellulo* by recruiting O-GlcNAc transferase (OGT), with temporal and magnitude control. Mass spectrometry data revealed that OGTACs induced site-selective O-GlcNAcylation of BRD4. Overall, OGTACs represent a promising approach for inducing protein-specific O-GlcNAcylation, thus enabling functional dissection and offering new directions for O-GlcNAc-targeting therapeutic development.

## INTRODUCTION

Protein O-linked β-*N*-acetylglucosamine modification (O-GlcNAcylation) plays a significant role in the regulation of transcription, metabolism, cell signaling, protein stability, and nucleocytoplasmic trafficking.^1–4^ The abnormal regulation of global O-GlcNAcylation state is associated with many human diseases, including cancer,^4–8^ neurodegenerative diseases,^9^ cardiovascular diseases,^10^ autoimmune diseases,^11^ and diabetes.^12^

However, the functional roles of O-GlcNAcylation are protein-specific, and the dissection of the functional consequences of individual O-GlcNAc modification events remains highly challenging.^2^ Current strategies used to study protein-specific O-GlcNAcylation include glycosite genetic mutation,^13,14^ dual-specific RNA aptamers,^15^ and nanobody-induced OGT-substrate proximity *in cellulo*,^16, 17^ which require extensive genetic engineering and are not amenable to magnitude control. Alternative techniques, such as protein semi-synthesis by peptide-protein ligation, are useful *in vitro* but difficult to apply in a cellular setting.^18^

Chemical inhibitors of O-GlcNAc transferase (OGT) and O-GlcNAcase (OGA), the enzymes that catalyze the addition and removal of O-GlcNAc, have also been intensively studied.^19^ However, as OGT and OGA are the only pair of enzymes regulating the O-GlcNAcylation of thousands of nuclear and cytosolic proteins, the inhibition of these enzymes inevitably disrupts the global cellular O-GlcNAcylation homeostasis, rendering the functional dissection and manipulation of protein-specific O-GlcNAcylation impossible. Moreover, prolonged treatment with these inhibitors can trigger feedback regulation of OGT/OGA, leading to confounding results from chemical perturbations.^20, 21^ Encouraged by the development of chemically induced proximity (CIP),^22^ we hypothesized that a heterobifunctional system that recruits OGT to the target protein could achieve target-specific O-GlcNAcylation. Unlike conventional OGA inhibitors, which directly block deglycosylation and thereby sustain high levels of O-GlcNAcylation, our heterobifunctional molecules are designed to bind OGT and a protein of interest (POI) simultaneously, inducing their proximity and triggering POI-specific O-GlcNAcylation.

Here, we present the development of O-GlcNAcylation targeting chimeras (OGTACs) which selectively induce O-GlcNAcylation of (theoretically) any POI, as demonstrated here by BRD4, CK2α, and EZH2, through promoting proximity with OGT. We optimize the molecular structure of OGTACs, and validate that their O-GlcNAcylation-inducing effect is dose- and time-dependent. Furthermore, through protein LC-MS/MS, we characterize previously unknown O-GlcNAcylation sites on BRD4.

## RESULTS AND DISSCUSSION

### Design of general OGTACs to induce targeted protein O-GlcNAcylation in cells

To demonstrate the feasibility of inducing protein-specific O-GlcNAcylation by chemically induced proximity, we first established a general, tag-based system for a proof-of-concept study. Since OGT activity is required, and there are currently no non-inhibitory ligands available for OGT, we exploited FKBP12^F36V^ fused-OGT (fOGT) as the O-GlcNAc transferase, which can be efficiently recruited by the AP1867 motif. Halotag-fused POIs were overexpressed as O-GlcNAcylation targets, which can be covalently captured by haloalkanes.^23^ We synthesized a series of molecules linking AP1867 and a haloalkane through different repeats of polyethylene glycol (PEG), producing O-GlcNAcylation-Targeting Chimeras (OGTACs) (**Figure 1a, b**). These OGTACs are designed to induce proximity between fOGT and any Halotag-fused POIs while concurrently promoting O-GlcNAc transfer to these specific POIs.

**Figure 1.**
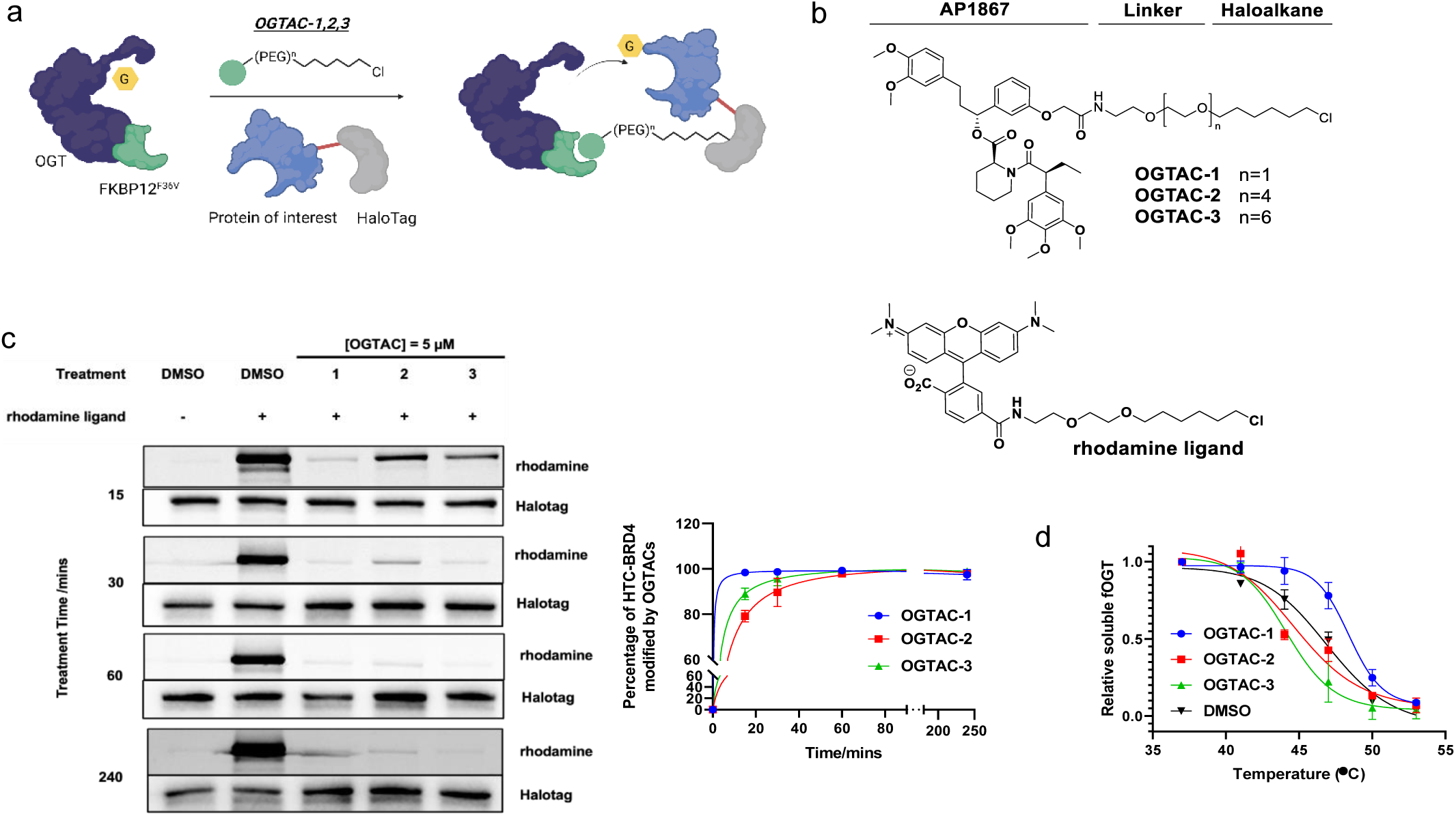
The general concept of OGTAC technology and target engagement studies of OGTAC molecules. (a), Conceptual scheme of OGTACs, which induce POI-specific O-GlcNAcylation by recruiting FKBP12^F36V^-OGT to Halotag-fused POIs. (b) Chemical structure of OGTAC-1,2,3, and rhodamine ligand for the pulse-chase experiment. (c, d) Confirmation of OGTAC1,2,3 target engagement in cells. HEK293T cells were transiently transfected with plasmids HTC-BRD4:fOGT at a 20:1 ratio. (c) In the pulse-chase assay, OGTAC-1,2,3 (5 μM) or vehicle were added to cells for increasing periods of time, followed by replacement with rhodamine ligand to label any remaining HTC-BRD4 proteins that OGTACs had not engaged. Then, cells were lysed and submitted to SDS-PAGE for in-gel fluorescence in the rhodamine channel and immunoblotted using a Halotag antibody to verify equal loading. (d) After 4 h treatment of 5 μM OGTAC-1,2,3, intact cell CETSA was conducted to verify their direct binding to fOGT. Data in (c, d) represent mean ± s.d. of n = 3 replicates.

### OGTAC-1 induces BRD4 O-GlcNAcylation in a dose and time-dependent manner

We selected bromodomain-containing protein 4 (BRD4) as the first POI for our proof-of-concept study. BRD4 is an epigenetic regulator that plays a significant role in cancer.^24^ Post-translational modifications (PTMs) of BRD4, such as phosphorylation, methylation, and ubiquitination, are well studied; phosphorylation was reported to affect chromatin targeting, onco-factors recruitment, and cancer progression.^25^ However, O-GlcNAcylation of BRD4 has never been studied comprehensively due to the lack of appropriate tools.

We first constructed the Halotag-C-terminal BRD4 (HTC-BRD4) plasmid and co-transfected it with fOGT plasmids in HEK293T cells at different ratios from 1:1 to 50:1. By immunoprecipitation (IP) and probing for O-GlcNAc using RL2 antibody in western blot (IP-WB), we observed that the O-GlcNAcylation level of HTC-BRD4 was proportional to the amount of fOGT expressed and only commenced when fOGT expression reached a threshold value (**Figure S1**). We, therefore, set the co-transfection ratio at 20:1, the threshold that showed minimal O-GlcNAcylation, for our OGTAC induction study.

Before evaluating the inducing effects of OGTACs, two cell-based assays were established to verify their ability to engage HTC-BRD4 and fOGT in cells. As the OGTAC-1/2/3 were designed to be irreversibly linked to the Halotag, we developed a pulse-chase assay to assess their HTC-BRD4 engagement efficacy. We first treated HEK293T cells co-expressing HTC-BRD4 and fOGT with OGTACs for different periods; then, we replaced the culture media and exposed the cells to a rhodamine ligand, which fluorescently labeled the HTC-BRD4 that did not interact with OGTACs (**Figure 1c**). Four time points were set for the pulse of OGTAC treatment, and the rhodamine signal on HTC-BRD4 was evaluated by in-gel fluorescence; low rhodamine signal indicated that the OGTACs had already reacted with the Halotag and the binding of the rhodamine ligand was blocked. All three OGTACs efficiently engaged HTC-BRD4, almost fully labeling the protein within 4 h. OGTAC-1 was the most efficient among the three OGTACs, achieving > 90% linkage within 15 mins of treatment.

For fOGT engagement, we established a cellular thermal shift assay (CETSA). Using the same co-expression system, we treated cells with OGTAC-1/2/3 for 4 h and conducted a heat challenge to the intact cells at a series of temperatures to determine the corresponding melting temperature (T_m_). OGTAC-1 treated groups showed increased T_m_ compared with the DMSO control with ΔT_m_ = 1.4 ℃ (**Figure 1d**), indicating that OGTAC-1 bound the fOGT protein and thereby stabilized the protein. However, OGTAC-2/3 showed no stabilizing effect on fOGT. As the principle of CETSA is to quantify the relative soluble protein after heat shock, we suspect that OGTAC-2 and 3 did not increase T_m_ due to the stable ternary complex they induced (shown in the next section, **Figure 2a**). The large ternary complex (molecular weight ∼ 400 kDa) is more likely to become insoluble upon heat challenge above 41 ℃.

**Figure 2.**
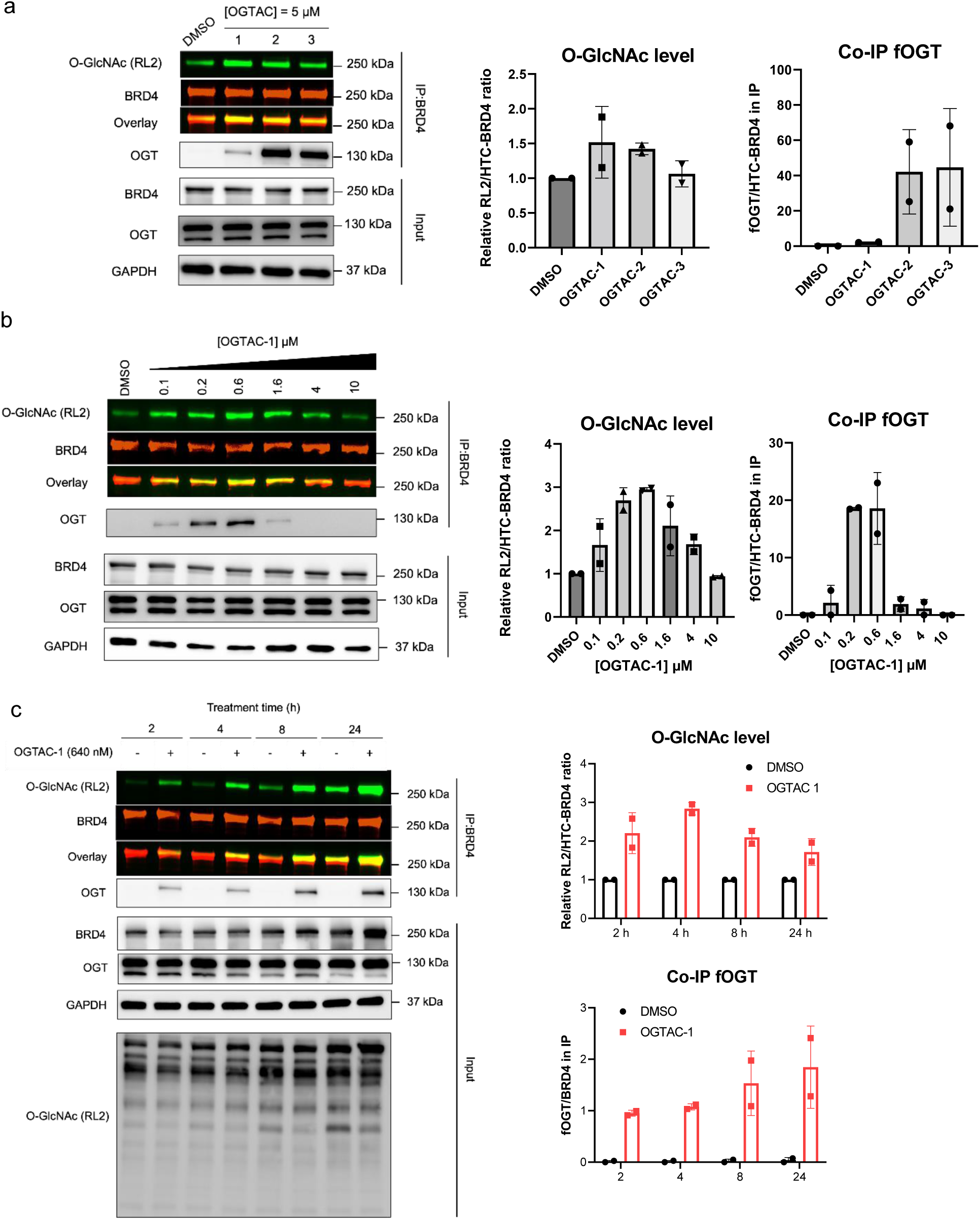
OGTAC-1 induces dose-dependent, rapid, targeted O-GlcNAcylation of HTC-BRD4 by recruiting fOGT in cells. (a), OGTAC-1 showed the best O-GlcNAcylation-inducing potency among three OGTACs at 5 μM. (b), Dose-dependent O-GlcNAcylation profile of HTC-BRD4 by OGTAC-1 treatment. HEK293T cells expressing fOGT:HTC-BRD4= 1:20 were treated with increasing concentration of OGTAC-1 for 4 h. Then, the O-GlcNAcylation level of HTC-BRD4 was assessed by immunoblot after IP using BRD4 antibody. Co-IP of fOGT was also observed in an OGTAC-1 dose-dependent manner. The right panel shows the quantitation of the immunoblot signal of RL2 relative to HTC-BRD4 as the mean ± s.e.m. of n = 2 biologically independent experiments.

We next assessed the induction of HTC-BRD4 O-GlcNAcylation in cells. We treated HEK293T cells co-transfected with HTC-BRD4 and fOGT with OGTAC-1/2/3 for 4 h, followed by immunoprecipitation of BRD4 proteins from the cell lysate and measurement of the O-GlcNAcylation level by immunoblot with RL2 antibody. All three OGTACs induced O-GlcNAcylation on HTC-BRD4, with OGTAC-1 showing highest potency (**Figure 2a**), consistent with the binding assays.

We further investigated the dose-dependent effect of OGTAC-1. The inducing effect was bell-shaped, with the highest level of O-GlcNAcylation observed between 256 nM and 640 nM. Co-IP of the fOGT was also observed at these concentrations, indicating that the induction of HTC-BRD4 O-GlcNAcylation depends on stable ternary complex formation (**Figure 2b**). The O-GlcNAcylation-inducing effect of OGTAC-1 declined from 1.6 μM, demonstrating that a hook effect occurred at this concentration. Due to the low expression level of endogenous BRD4, we could not observe the effects of OGTAC-1 on it using immunoblot.

Following the identification of the optimized concentration of OGTAC-1, we evaluated the kinetics of OGTACs-mediated O-GlcNAcylation to reveal time-dependent effects. We examined the O-GlcNAcylation level at 2 h, 4 h, 8 h, and 24 h. The RL2 signal of HTC-BRD4 displayed a 2.5-fold increase at 2 h and further escalated to a 3-fold increase at 4 h (**Figure 2c**). However, longer treatment did not further enhance O-GlcNAcylation, possibly due to the limitations of the transient transfection system: although media with transfection reagents and plasmids were removed before OGTAC treatment, the cells were still expressing exogenous HTC-BRD4 and fOGT. The ratio of fOGT:OGTAC:HTC-BRD4 we optimized for 4 h treatment was compromised, leading to weaker potency of the inducing effect. Importantly, we did not observe increased O-GlcNAcylation on other proteins (**Figure 2c**), confirming the target-specific inductive effect of OGTAC-1.

Collectively, these data suggest that OGTAC-1 induced BRD4 O-GlcNAcylation in a dose-dependent and time-dependent manner, without altering the global O-GlcNAcylation level.

### OGTAC strategy generally applied to other POIs

Next, to illustrate the generality of our OGTAC strategy, we assessed OGTAC-1/2/3 effects on HTC-Casein kinase (HTC-CK2α). CK2α is a well-studied OGT substrate.^18, 26^ Ser347 is conservatively O-GlcNAcylated, and this modification was reported to affect its kinase activity.^17, 18^ We first co-expressed HTC-CK2α and fOGT in HEK293T cells at different ratios and found that the basal O-GlcNAcylation level of HTC-CK2α was higher than what we detected in the case of HTC-BRD4. Therefore, a lower ratio of fOGT expression (1:100) was selected to assess OGTAC potency (**Figure S2**). We first conducted IP-WB, and the result indicated that OGTAC-1 was again the most efficient among the three OGTACs, although it did not yield co-IP quantifiable fOGT (**Figure 3a**). The induction of specific O-GlcNAcylation on HTC-CK2α was further confirmed by a mass-shift assay. In this assay, O-GlcNAcylated proteins in cell lysate were enzymatically labeled with N-azidoacetylgalactosamine (GalNAz), followed by covalent linkage to polyethylene glycol (PEG) of 5 kDa by strain-promoted alkyne-azide cycloaddition (SPAAC), resulting in an up-shifted band in immunoblot (**Figure 3b**).^27^ We validated that 1 μM of OGTAC-1 increased HTC-CK2α O-GlcNAcylation by 1.5 fold; importantly, OGTAC-1 treatment did not affect the endogenous CK2α O-GlcNAcylation level, suggesting its target specificity (**Figure 3b**).

**Figure 3.**
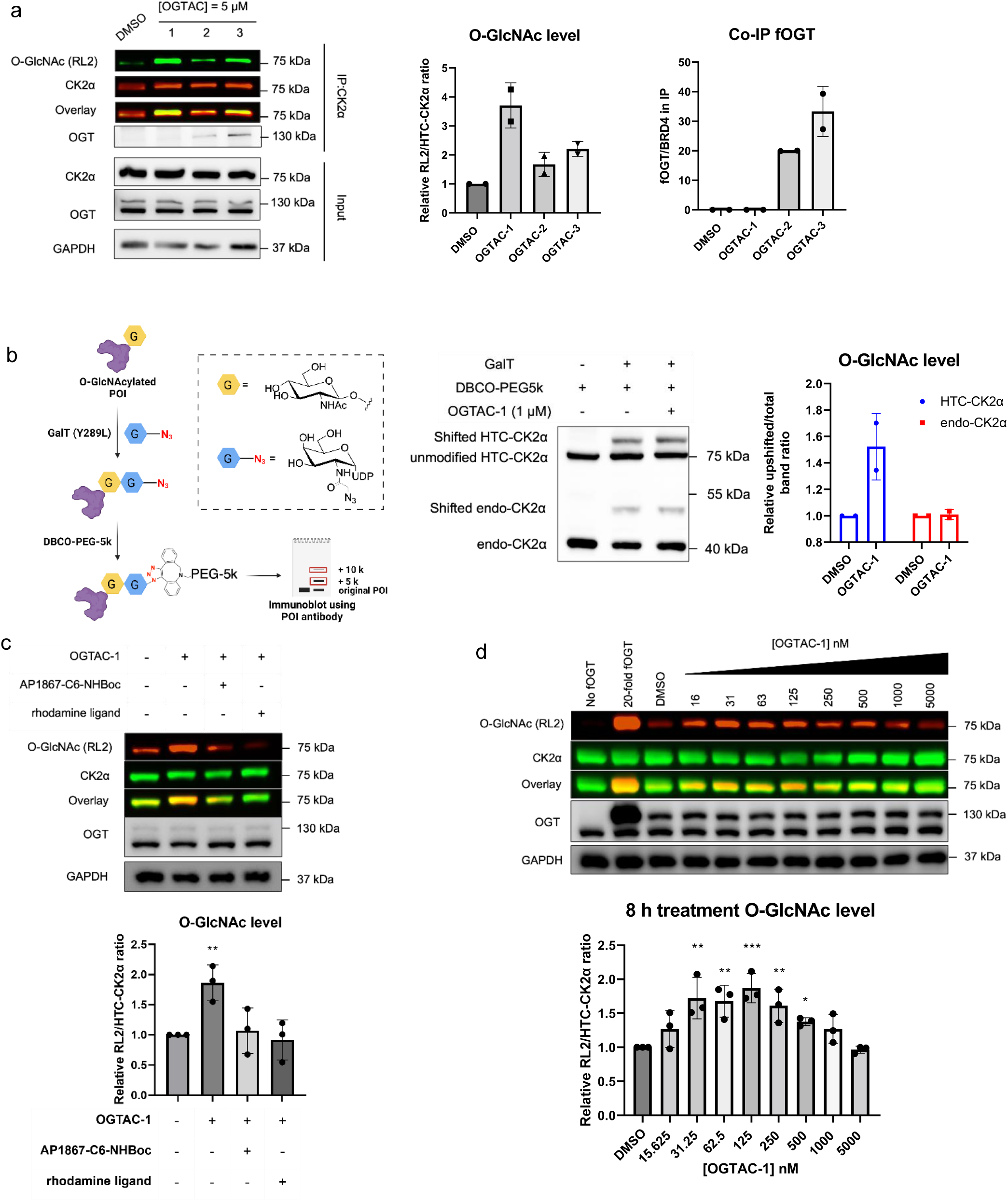
OGTAC technology extended to HTC-CK2α. (a), IP-WB method showed OGTAC induced HTC-CK2α O-GlcNAcylation; (b), Mass-shift assay to validate the OGTAC-1 induced specific HTC-CK2α O-GlcNAcylation; (c), immunoblot analysis of OGTAC-1 mediated O-GlcNAcylation when co-treated with binding competitors; (d), OGTAC-1 induced dose-dependent, time-dependent (**Figure S5,6**) O-GlcNAcylation of HTC-CK2α in cells. HEK293T cells were transfected with fOGT:HTC-CK2α=1:100 for 24 h, followed by treatment with increasing concentration of OGTAC-1 for 8 h. All quantifications are shown as mean ± s.e.m. of 3 independent biological repeats. Statistical significance in (D) was calculated with unpaired two-tailed Student’s t tests comparing DMSO-to OGTAC-1-treated samples. *p < 0.05; **p < 0.01; ***p < 0.001.

During the immunoblotting study, we noticed that even in whole cell lysate (WCL), the O-GlcNAcylation on HTC-CK2α was quantifiable (**Figure S3**). We assessed the OGTAC treatment effects by quantifying the RL2 signal overlapping with HTC-CK2α in WCL and found that they corresponded with the results achieved by the IP-WB method (**Figure S4**). Therefore, in later studies, we directly quantified the intensity of the RL2 band overlapping with HTC-CK2α instead of IP HTC-CK2α. To validate that direct binding to both fOGT and HTC-CK2α is critical, we exploited AP1867 and rhodamine ligand, which bind to fOGT and HTC-CK2α, respectively, and thus compete with OGTAC-1. As expected, co-treatment with either competitor abolished the O-GlcNAcylation-inducing effects of OGTAC-1 (**Figure 3c**).

Next, we investigated the dose- and time-dependent effects of OGTAC-1. We observed that, at 250 nM, induction of O-GlcNAcylation occurred within 4 h, peaked at 8 h, and then started to decline (**Figure 3d, S5**). However, for 24 h treatment (**Figure S6**), a higher concentration of OGTAC-1 (1 μM) demonstrated significant effects; this is because a greater concentration of OGTAC-1 is necessary to sustain the appropriate stoichiometry with the increased expression of HTC-CK2α protein. This phenomenon again indicates the importance of optimizing fOGT:OGTAC:POI ratios for efficient chemically induced O-GlcNAcylation.

The same strategy was also extended to enhancer of zeste homolog 2 (EZH2), an epigenetic regulator that is an OGT substrate.^28, 29^ OGTAC-1 also demonstrated the highest O-GlcNAcylation-inducing effects among the three OGTACs (**Figure S7**). As there are >20,000 commercially available plasmids for Halotag-fused human proteins,^30^ OGTACs can be rapidly tested in a wide range of POIs, enabling time- and dose-dependent manipulation of their specific O-GlcNAcylation level.

### OGTAC-4 induces BRD4-specific O-GlcNAcylation

The successful application of OGTAC-1 demonstrated the viability and generality of chemically induced O-GlcNAcylation in a dual fusion-protein system. We next pursued the induction of POI-specific O-GlcNAcylation by direct binding to the native domain of POIs; this could facilitate the induction of O-GlcNAcylation on endogenous POIs for functional dissection. To this end, we synthesized OGTAC-4, in which a JQ1^31^ structure was incorporated as a binding motif for the BRD4 BD1/BD2 domain (**Figure 4a, b**). As OGTAC-4 directly recruits BRD4, we first tried to assess its potency on endogenous BRD4 in a cell line stably expressing fOGT. However, no O-GlcNAcylation signal for endogenous BRD4 could be detected by the IP-WB method using RL2 as the antibody (data not shown). We therefore overexpressed BRD4 together with fOGT in HEK293T cells to accommodate the readout within the IP-WB detection limit. In this co-expression system, we validated the O-GlcNAcylation-inducing effect of OGTAC-4 **(Figure 4c)**. Of note, OGTAC-4 promoted BRD4 O-GlcNAcylation and the formation of a stable ternary complex between BRD4 and fOGT at concentrations as low as 16 nM (**Figure 4c & S8**). Compared with the covalent inducer OGTAC-1, OGTAC-4 demonstrated a wider effective concentration range. This may be due to the sub-stoichiometric catalytic activity of OGTAC-4: at a lower concentration, OGTAC-4 can still be effective because, after O-GlcNAc transfer from fOGT to BRD4, the BRD4 would dissociate from the original ternary complex and unmodified BRD4 could be recruited to fOGT by OGTAC-4. The kinetics of OGTAC-4-induced O-GlcNAcylation correspond with this interpretation (**Figure 4d**). OGTAC-4 induced O-GlcNAcylation within 2 h, and the induction increased with time. Collectively, our results indicate that OGTAC-4 has catalytic characteristics and can be applied at low concentration with long-lasting effects.

**Figure 4.**
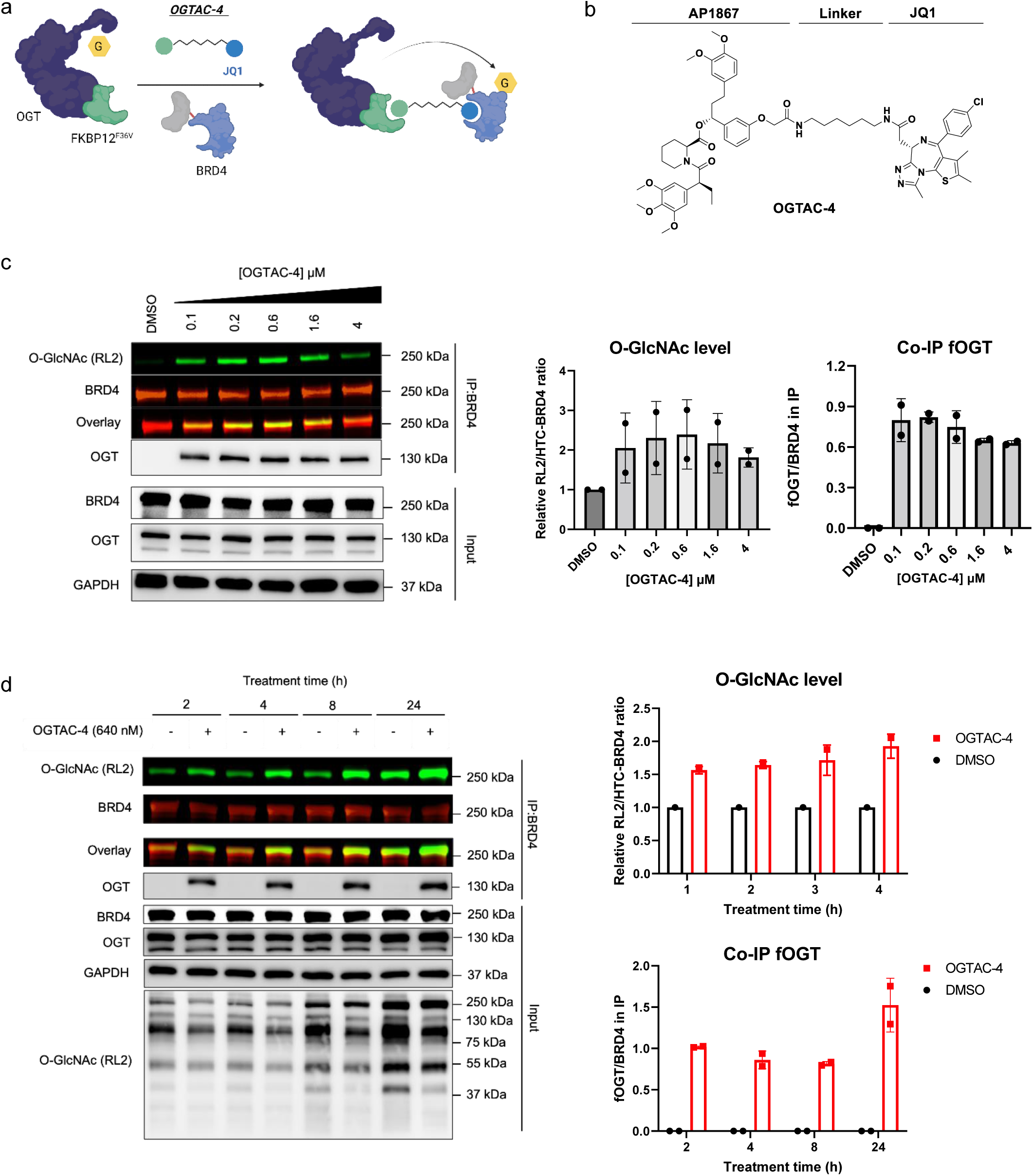
OGTAC-4 induces BRD4 O-GlcNAcylation by recruiting fOGT in cells; (b), Chemical structure of OGTAC-4; (c), Dose-dependent O-GlcNAcylation profile of HTC-BRD4 by OGTAC-4 treatment. HEK293T cells expressing fOGT:HTC-BRD4= 1:20 were treated with an increasing concentration of OGTAC-4 for 4 h. Then, the O-GlcNAcylation level of HTC-BRD4 was assessed by immunoblot after IP using BRD4 antibody. Co-IP of fOGT was also observed in an OGTAC-4 dose-dependent manner; (d), OGTAC-4 (640 nM) O-GlcNAcylation-inducing effects on HTC-BRD4 over indicated time course. The right panel shows the quantitation of the immunoblot signal of RL2 relative to HTC-BRD4 as the mean ± s.e.m. of n = 2 biologically independent experiments

### OGTACs mediate site-selective O-GlcNAcylation of BRD4

Although both OGTAC-1 and OGTAC-4 effectively induced POI-specific O-GlcNAcylation, the modified glycosites remain unknown. To elucidate the site selectivity of O-GlcNAcylation induced by the two OGTACs, we mapped the modified glycosites of BRD4 using liquid chromatography-coupled tandem mass spectrometry (LC-MS/MS). In these experiments, we transfected HEK293T cells with HTC-BRD4:fOGT = 20:1 for 24 h, then treated cells with 1) DMSO, 2) OGTAC-1, or 3) OGTAC-4 for 4 h. HTC-BRD4 proteins were enriched by HaloTrap agarose instead of BRD4 antibody for better elimination of the background signal from endogenous BRD4. Considering the low stoichiometry of O-GlcNAcylation, we further separated the eluates by sodium dodecyl sulfate–polyacrylamide gel electrophoresis (SDS-PAGE) and cut out the bands for HTC-BRD4 to maximize the detection of all O-GlcNAcylated peptides with LC-MS/MS (**Figure 5a**). In parallel with MS analysis, we also used immunoblotting to verify the induction of HTC-BRD4 O-GlcNAcylation and its specificity (**Figure S9**). To avoid random or artificial effects from the testing and database search, only O-GlcNAcylation sites detected in at least two of three biological repeats were used for quantification. The MS results indicated that both OGTAC-1 and OGTAC-4 induced neo-O-GlcNAcylation at T298, T299, S1212, and S1262 (**Figure 5b**) (unmodified in DMSO group). Interestingly, OGTAC-4 induced unique S1117 O-GlcNAcylation, which was undetected both in DMSO and OGTAC-1 treatment. Among the 24 consensus O-GlcNAc sites (**Figure 5b**), quantitative proteomics revealed that O-GlcNAcylation of S1122 was enhanced by both OGTAC-1 and OGTAC-4, while that of S1051 was exclusively enhanced by OGTAC-1 (**Figure 5c**). These data collectively show that OGTAC-1 and OGTAC-4 induced site-selective BRD4 O-GlcNAcylation, which can facilitate the site-specific dissection of BRD4 O-GlcNAcylation.

**Figure 5.**
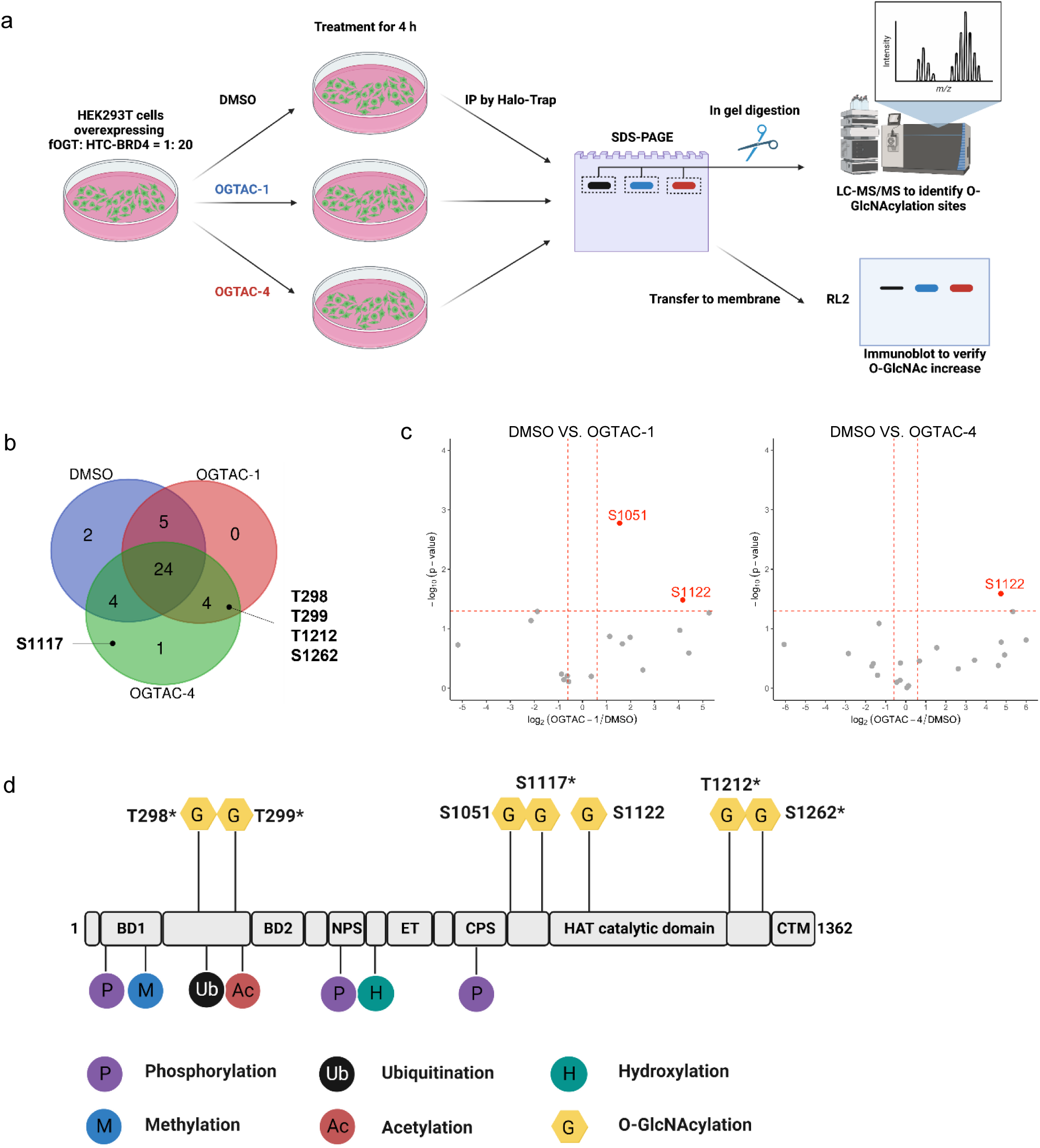
Quantitative proteomics to validate induction of BRD4 O-GlcNAcylation. (a), sample preparation for the O-GlcNAcylation site identification by LC-MS/MS. Cells treated with DMSO, OGTAC-1 (640 nM), or OGTAC-4 (640 nM) for 4 h were lysed, enriched by Halo-Trap, eluted and separated by SDS-PAGE, the gel bands for HTC-BRD4 were cut, trypsinized for quantitative MS. The same eluted protein samples were submitted to immunoblot to verify the treatment effects (**Figure S8**); (b), Venn diagram showing identified O-GlcNAc sites from DMSO/OGTAC-1/OGTAC-4 treatment. Neo-sites discovered after probe treatment are labeled next to the corresponding area. O-GlcNAc sites only detected in one repeat were ignored from quantifications; (c), Volcano blots showing OGTAC-1 and 4 induced site-specific O-GlcNAcylation. Peptides were prepared according to the method described in (a) and then subjected to quantitative MS. The vertical dashed lines correspond to a 1.5-fold change relative to DMSO, and the horizontal line corresponds to a P value of 0.05 for statistical significance. Red points correspond to sites with >1.5-fold change (P < 0.05) relative to DMSO. Each point represents an individual O-GlcNAcylated site plotted as the mean of n = 3 biological replicates. O-GlcNAc sites only detected in one repeat were ignored from quantifications;(d), Reported PTMs and novel O-GlcNAcylation sites induced by OGTACs on full-length BRD4. Asterisk indicates a neo-O-GlcNAc site induced by OGTACs. BRD4 domains that have reported functions are labeled with short forms as follows: NPS, N-terminal phosphorylation site; ET, extra-terminal domain; CPS, C-terminal phosphorylation site; HAT, Histone acetyltransferases; CTM, C-terminal domain.

By comparing the loci of all induced O-GlcNAc sites with other BRD4 residues with reported PTMs, we identified sites such as T298 and T299 O-GlcNAcylation that potentially engage in crosstalk with ubiquitination,^32^ whereas S1122 may influence its histone acetyltransferase activity (**Figure 5d**).^33^ Since the BRD4 C-terminus interacts with and destabilizes the protooncogene MYC, O-GlcNAcylation on S1117, T1212, and S1262 could potentially interfere with this cancer-associated interaction.^34^ Our results could catalyze follow-up experiments (e.g. site-directed mutagenesis studies) to validate these putative functions of site-specific O-GlcNAcylation.

## CONCLUSION

Here, we developed the first small molecule-based technology, OGTAC, for protein-specific induction of O-GlcNAcylation in living cells. OGTAC-1 is a chemical inducer of O-GlcNAcylation generally applicable for tagged POIs, as shown here by BRD4, CK2α, and EZH2. In addition, OGTAC-4 was developed for induction of BRD4-specific O-GlcNAcylation. Both OGTAC-1 and OGTAC-4 are sub-micromolar efficacy O-GlcNAcylation inducers, promoting the formation of stable ternary complex. OGTAC-1 covalently bound the POIs and demonstrated autoinhibition of O-GlcNAcylation at higher concentrations, while OGTAC-4 functioned as a sub-stoichiometric catalyst for BRD4-specific O-GlcNAcylation. Furthermore, LC-MS/MS revealed that their inducing effects on BRD4 are site-selective, which can facilitate site-specific functional elucidation for BRD4 O-GlcNAcylation, in concert with other techniques (e.g. site-directed mutagenesis).

Despite the growing recognition of the significant roles that protein-specific O-GlcNAcylation plays in various cellular processes and diseases, the methods to uncover these roles are still underdeveloped. Genetic mutation of glycosites^28, 35, 36^ can help elucidate the site-specific function of O-GlcNAcylation. However, its limitations are inevitable: 1) mutation can affect protein folding, substrate recognition, and competing PTMs such as phosphorylation. 2) as a dynamic and sub-stoichiometric modification, temporal and magnitude control of O-GlcNAcylation is crucial but cannot be achieved. Although nanobody-OGT^16^ and dual-specific RNA aptamers^15^ have been recently developed, their mechanisms still constrain them from precise temporal and magnitude control. Our OGTAC technology, as a small molecule-based method, enables general, rapid, dose-dependent POI-specific O-GlcNAcylation.

During the development of OGTAC technology, we noticed that linker length is critical for optimized O-GlcNAc transfer from OGT to POIs. According to target engagement assays and the evaluation of O-GlcNAcylation-inducing effects, OGTAC-2 and 3 promote the formation of stable ternary complexes but show poor O-GlcNAcylation-inducing effects. We conclude that longer linkers do not favor the O-GlcNAc transfer from OGT to POIs. We rationalize this as follows: 1) Our OGTACs bind to both proteins with high affinity. For OGTAC-1/2/3, the AP1867 motif binds to FKBP12^F36V^ with a K_d_= 1.8 nM; the other terminus covalently binds to Halotag-POI. These hyper-potent binders induce the proximity between fOGT and Halotag-POIs and lock their relative position. The distance between the proteins thus depends on the OGTAC linker length. A greater distance between the two proteins caused by the longer linker could hamper O-GlcNAc transfer from the fOGT catalytic site to Halotag-POIs. Although the longer linker reduces the potential clash between fOGT and POIs, promoting the formation of a stable ternary complex, a closer distance between the catalytic site of fOGT and POIs or even direct binding between fOGT and POIs is more crucial. In most cases, OGT needs to directly recognize the substrate with its tetratricopeptide repeat (TPR) domain, highlighting the significance of the physical distance between OGT and its substrate.^37, 38^ It has been reported that protein substrates with O-GlcNAc sites around their C-terminus may bypass this TPR domain binding.^38^ We suspect that the glycosites we detected that are proximal to the BRD4 C-terminal domain (CTM), such as S1212 and S1262, are the most accessible sites for O-GlcNAc transfer and may be independent of TPR recognition. These sites may also be induced by OGTAC-2/3, while the potency gap between OGTAC-1 and OGTAC-2/3 might be owing to the remaining O-GlcNAc sites, which require BRD4 interaction with the TPR domain of fOGT.

While both OGTAC-1 and OGTAC-4 are bifunctional small molecule inducers for BRD4 O-GlcNAcylation, they exhibit distinct characteristics. The most obvious differences lie between their working concentration and time-dependent activity. OGTAC-1 demonstrated an obvious autoinhibitory bell-shaped curve in the dose-dependent study. However, OGTAC-4 showed an effect below 100 nM, and no autoinhibitory concentration was found even at 10 μM (**Figure 4c & S8**). Considering AP1867 is a low nanomolar binder for FKBP12^F36V^ (K_d_ = 1.8 nM)^39^, the JQ1 motif of OGTAC-4 (K_d_ = 49.0 nM for BD1, 90.1 nM for BD2)^31^ is more likely to dissociate and recruit additional BRD4 as the substrate for the subsequent O-GlcNAc transfer reaction. In contrast, the covalent binding between OGTAC-1 and BRD4 inhibits this dissociation and reassociation process. As a result, OGTAC-4 but not OGTAC-1 can act as a catalyst for O-GlcNAc transfer. Consequently, prolonged treatment is favored for OGTAC-4 but not for OGTAC-1.

Although we successfully achieved targeted O-GlcNAcylation using bifunctional molecules, the current approach still has several limitations. The lack of a non-inhibitory ligand for OGT mandates exogenous expression of fOGT, which hinders the translation of these small molecules to therapeutic applications. The discovery of a binder for recruiting native OGT in cells would be an important step forward. Furthermore, methods to detect and quantify protein-specific O-GlcNAcylation still need to be improved, and quantitative proteomics is indispensable to accurately determine the O-GlcNAcylation status at the Ser/ Thr site level. In summary, our OGTAC technology demonstrates the potential of using small molecules to induce protein-specific O-GlcNAcylation. This breakthrough provides a novel tool for more deeply understanding the functional roles of protein O-GlcNAcylation. Additionally, it lays the foundation for therapeutically targeting dysregulated O-GlcNAcylation in human diseases.

## AUTHOR INFORMATION

### Author Contributions

B.W.M. and K.S.K. contributed equally to this work.

## ACKNOWLEDGMENTS

The authors acknowledge Ms. Sarah Ng from Department of Chemistry, CUHK for their NMR facilities. We thank Prof. Timothy Mitchison (Harvard Medical School) and Prof. John Karanicolas (Fox Chase Cancer Center), Prof. Jiang Xia (CUHK) for their scientific discussions. We thank Prof. Suzanne Walker (Harvard Medical School), Prof. Yangchao Chen (CUHK) and Prof. Siaw Shi Boon (CUHK) for sharing plasmids. We thank Dr. Tom Wright for the academic editing on the manuscript. B.W.-L.N. acknowledges funding support from CUHK (CU Medicine Faculty Innovation Award; FIA2020/A/03) and Croucher Foundation (Start-up Grant).

