## Supplementary Information for "Targeted Protein O-GlcNAcylation Using Bifunctional Small Molecules"

**Supplementary figures:**

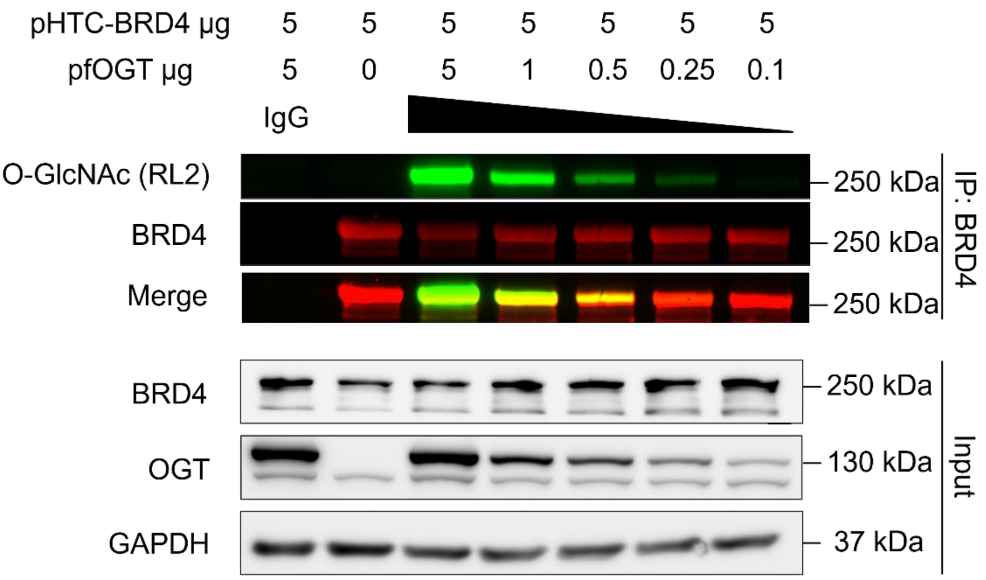


**Figure S1.** fOGT: HTC-BRD4 plasmid ratio optimisation. Co-transfection of pHTC-BRD4 5 µg with 0.25 µg (20:1) was selected for later study.

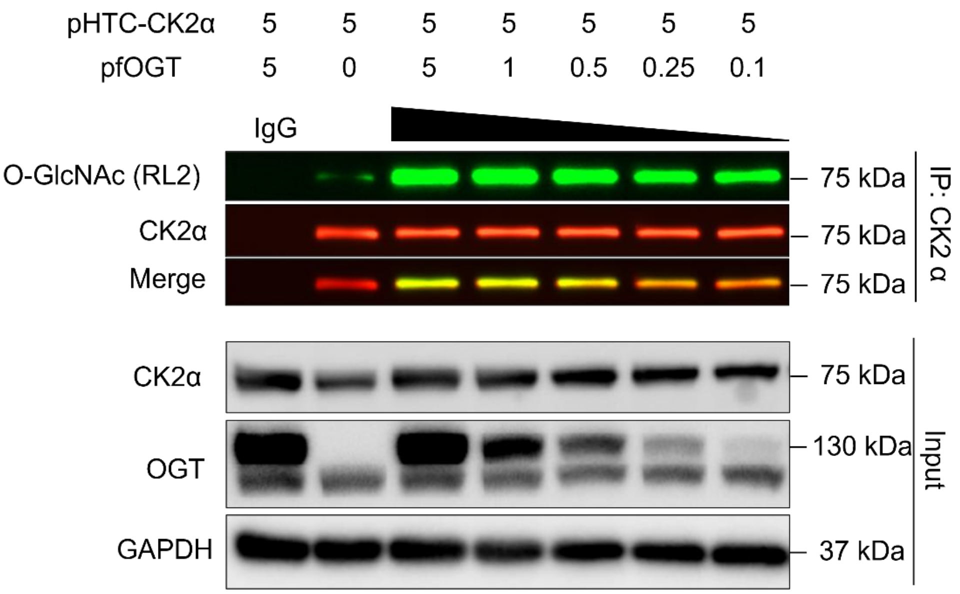


**Figure S2.** fOGT: HTC-CK2α plasmid ratio optimisation. Co-transfection of pHTC-CK2α 5 µg with 0.05 µg (100:1) was selected for later study.

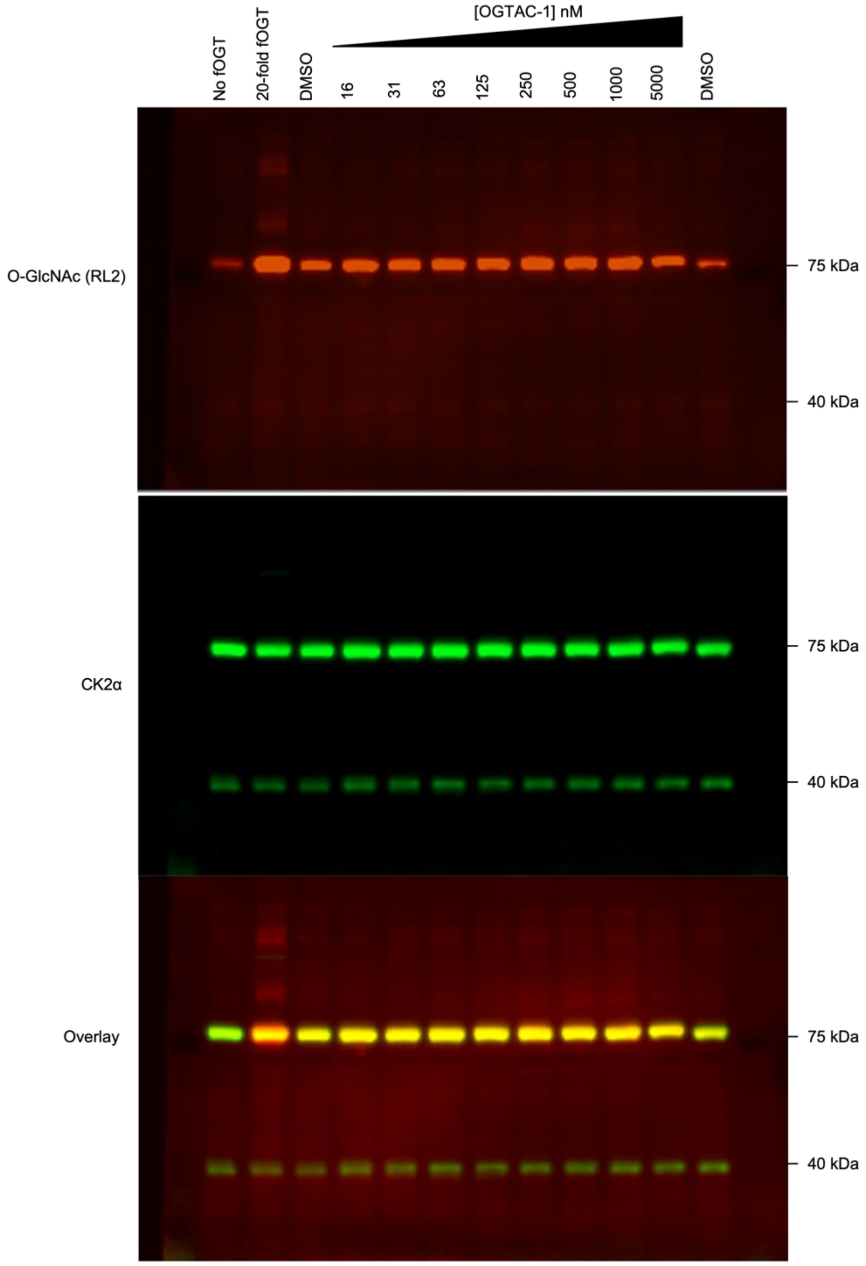


**Figure S3.** Evaluation of HTC-CK2α O-GlcNAc level from whole cell lysate WB. Overlapping the pan-RL2 (red) with HTC-CK2α (green at ~75 kDa) reveals the HTC-CK2α specific O-GlcNAc level.

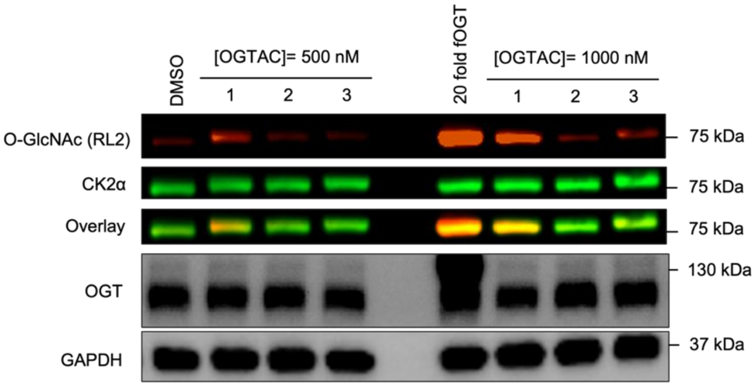


**Figure S4.** Evaluation of O-GlcNAc inducing effects of OGTAC-1/2/3 on HTC-CK2α from whole cell lysate WB. OGTAC-1 induced higher fold increase of O-GlcNAcylation on HTC-CK2α both at 500 nM and 1 µM

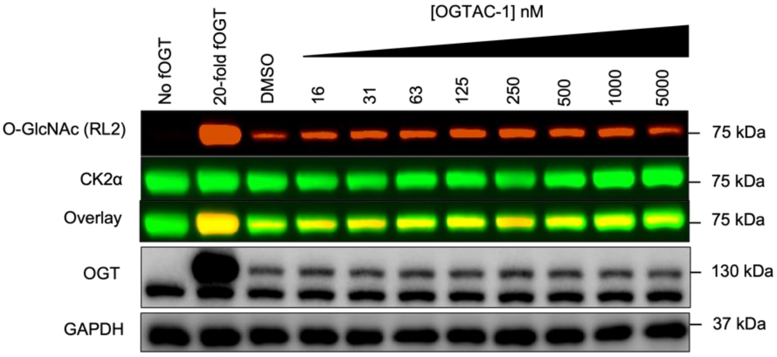


**Figure S5.** OGTAC-1 dose-dependent O-GlcNAc inducing effect on HTC-CK2α after 4 h treatment.

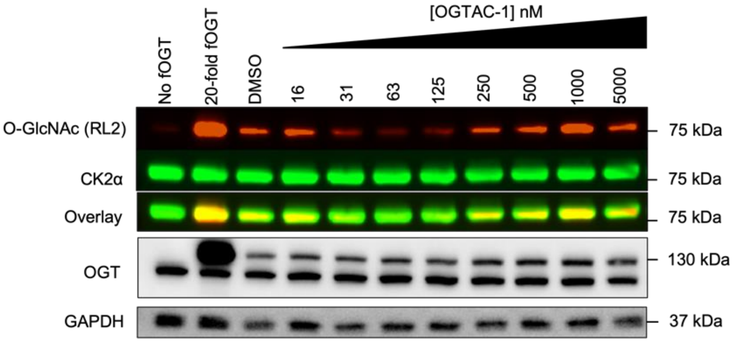


**Figure S6.** OGTAC-1 dose-dependent O-GlcNAc inducing effect on HTC-CK2α after 24 h treatment.

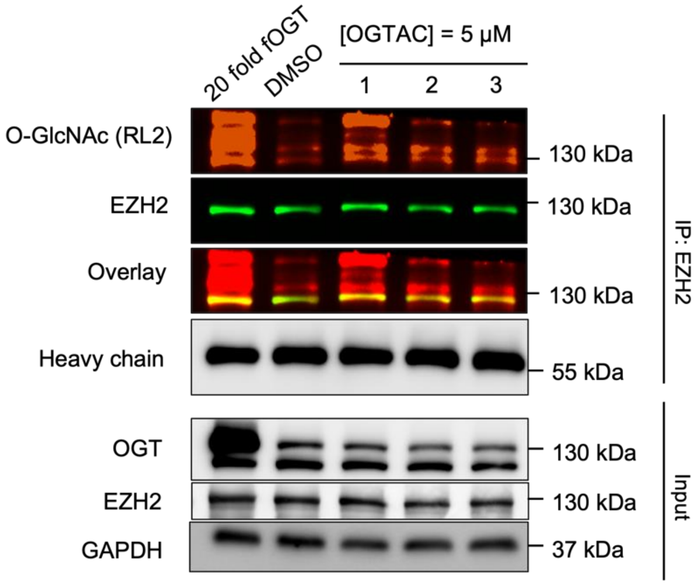


**Figure S7.** Evaluation of O-GlcNAc inducing effects of OGTAC-1/2/3 on HTC-EZH2 by IP-WB method. In the treatment group, pfOGT: pHTC-EZH2= 1:100.

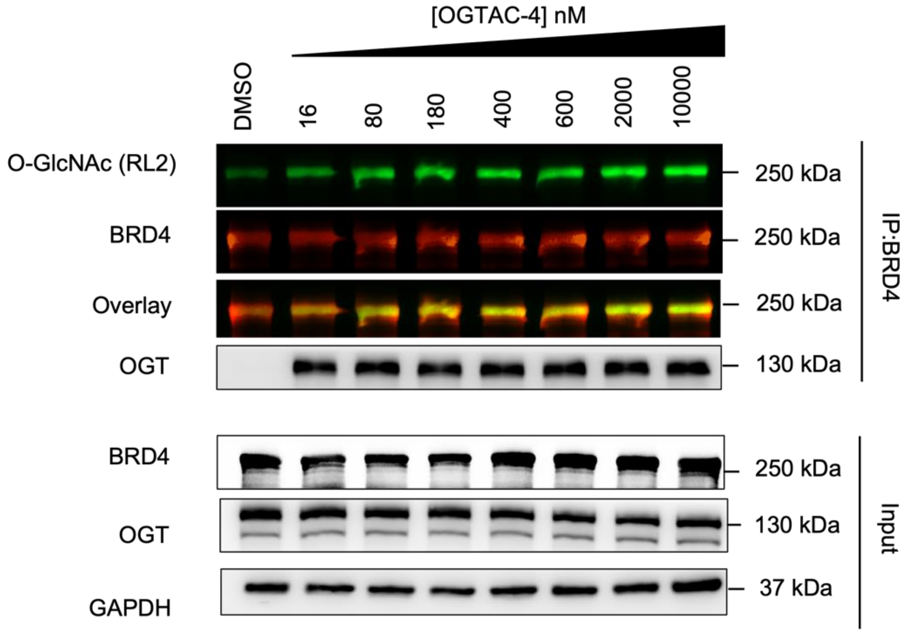


**Figure S8.** Evaluation of O-GlcNAc inducing effects at wider concentration range of OGTAC-4 on HTC-BRD4 by IP-WB method.

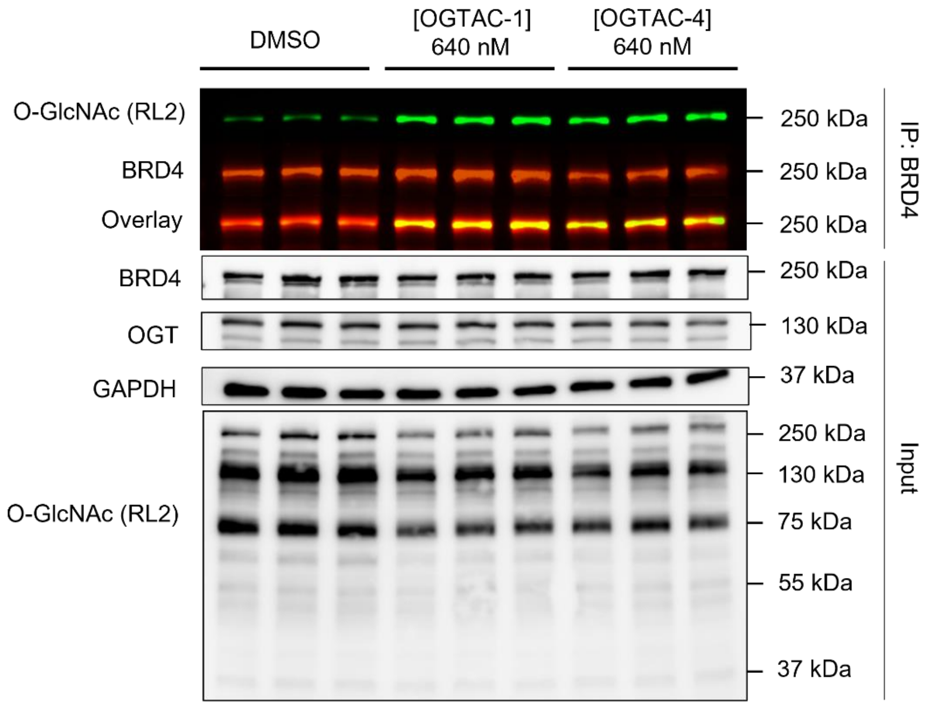


**Figure S9.** Evaluation of O-GlcNAc inducing effects of OGTAC-1 and OGTAC-4 on HTC-BRD4. Cells treated with DMSO, OGTAC-1 (640 nM), or OGTAC-4 (640 nM) for 4 h were lysed, enriched by Halo-Trap then submitted to WB.

### <u>Materials and Methods:</u>

#### Cell Cultures, DNA Constructs, and Reagents

HEK293T cells (kind gift from Prof. Sze Lok Cheng, CUHK) were maintained and cultured at 37 °C with 5% $CO_2$ in Dulbecco's modified Eagle's medium (DMEM) supplemented with 10 % fetal bovine serum and 1% penicillin and streptomycin.
FKBP12$^{F36V}$-OGT vector was a gift from Walker's group (Harvard Medical School).
HTC/HTN-BRD4 plasmids were constructed by subclone from pcDNA4-TO-HA-Brd4FL (addgene: #31351), then homologous recombination (Vazyme C115) into pHTC/N HaloTag® CMV-neo Vector (Promega G7711).
HTC/HTN-CK2α plasmids were constructed by subclone from pDB1 (CK2αlpha) (addgene: #27083), then homologous recombination into pHTC/N HaloTag® CMV-neo Vector.
HTC/HTN-EZH2 plasmids were constructed by subclone from 3XMyc-His$_6$-EZH2 plasmid from Prof. YangChao Chen (CUHK), then homologous recombination into pHTC/N HaloTag® CMV-neo Vector.

#### Pulse-chase assay

1 X 10$^5$ HEK293T cells were seeded in each well of 12 well plate. After cells attached, transfection reagents and plasmids were prepared and added in DMEM. After 24 h

transfection, media were replaced by fresh warm complete media with DMSO or compounds. After certain time of OGTACs treatment, media in wells were replaced by fresh warm media with 5 µM rhodamine ligand. The cells were incubated in 37 °C for 15 mins. Then, media were aspirated and 100 µL IP lysis buffer (Thermo Scientific 87788) supplemented with 20 µM OGA inhibitor (Thaimet G, Bidepharm BD571819) and 100X protease inhibitor (MedChemExpress, HY-K0010) (referred as complete IP lysis buffer in the later paragraph) was added to each well. The lysate was incubated on ice for 20 min, spun down at 14,000 RPM at 4°C for 15 mins, and the supernatant were collected for bicinchoninic acid (BCA) assay and normalised to a final 2 mg/mL. The lysate were denatured by SDS- loading buffer (Biorad #1610747) and 10 µL of each sample was loaded in SDS-PAGE. After running, in gel fluorescence at rhodamine channel (545/575 nm) was analysed by Bio-Rad ChemiDoc MP Imaging System. After image, the gel was transferred to PVDF membrane to check equal loading using anti-GAPDH antibody.

**CETSA**

HEK293T cells (2.5 X $10^6$ cells) were seeded in 10 cm culture dish overnight for attachment. On the next day, cells were transfected with 20:1 HTC-BRD4:fOGT plasmid for 24 hours until ~90% confluence. After treatment with 5 µM compounds or DMSO, cells were collected and washed with PBS. Subsequently, cells were re-suspended in 500 µL PBS and equally divided into 50 µL aliquots. The tubes were subjected to heat challenge at 37-53°C for 5 mins, followed by cooling on ice. Cells were lysed by three repeated freeze-thaw cycles with liquid nitrogen and water bath. Lastly, cells were centrifuged at 14000 RPM for 15 mins and supernatants were collected for Western Blot analysis.

**Immunoprecipitation**

For each IP reaction, 2.2 X $10^6$ HEK293T cells were seed in 10 cm dish. After cells attached, transfection reagents and plasmids were prepared and added in DMEM. After 24 h transfection, media were replaced by fresh warm complete media with DMSO or probes. After certain time of OGTACs treatment, cells were collected by cold PBS and lysed by 200 µL complete IP lysis buffer. The lysate was incubated on ice for 20 min, spun down at 14,000 RPM at 4°C for 15 mins, and the supernatant were collected for bicinchoninic acid (BCA) assay and normalised to a final 2 mg/mL. For input, 20 µL of diluted sample was reacted with equal volume of 10 µM TMR ligand in IP lysis buffer and rotate gently at RT for 15 mins. The remaining samples were subjected to protein A/G magnetic beads (MedChemExpress, HY-K0202), which pre-binding with protein target protein antibody. After gentle rotation at 4 °C for at least 16 h, the beads for wash with IP lysis buffer and boiled in 40 µL 2X SDS-loading buffer (Biorad #1610747) for 5 mins to elute proteins from beads. Eluted samples were directly subjected into WB analysis. To get stronger signal of RL2, we used anti-mouse-HRP (Cell signaling, #7076) as secondary antibody for RL2 and anti-rabbit-Alexa Fluor™ 488 (Invitrogen, #A-11008) as secondary antibody for total target proteins.

### Western blotting and Antibody

In general, cells (12 well) were lysed by 80 µl RIPA lysis buffer (Thermo Scientific 89901) supplemented with 20 µM OGA inhibitor (Thaimet G, Bidepharm BD571819) and 100X protease inhibitor (MedChemExpress, HY-K0010), incubated on ice for 20 min, spun down at 14,000 RPM at 4°C for 15 mins, and the supernatant were collected for bicinchoninic acid (BCA) assay and normalised to a final 2 mg/mL concentration. About 30 µg of protein samples were loaded for sodium dodecyl sulfatepolyacrylamide gel electrophoresis (SDS-PADE) and blotted with indicated antibodies. Antibodies used in this study are as follow:

| Antibody | Brand | Dilution (application) |
|---|---|---|
| RL2 | Abcam (ab2739) | 1:1000 (WB) |
| Anti-BRD4 | Cell Signalling Technology (CST) (13440S) | 1:2000 (WB)<br>1:100 (IP) |
| Anti-OGT | CST (D1D8Q) | 1:1000 (WB) |
| Anti-Ck2α | CST (2656) | 1:1000 (WB) |
| Anti-EZH2 | CST (5246) | 1:2000 (WB) |
| Anti-Ck2α | Proteintech (10992-1-AP) | 1:50 (IP) |
| Anti-GAPDH | Santa Cruz (sc-47724) | 1:3000 (WB) |
| Anti-Halotag | Promega (G9211) | 1:1000 (WB) |
| Goat anti-Mouse-HRP | CST (7076) | 1: 6000 (WB) |
| Goat anti-rabbit-HRP | CST (7074) | 1: 6000 (WB) |
| Goat anti-rabbit-Alexa Fluor™ 488 | Invitrogen (A-11008) | 1: 6000 (WB) |
| IRDye® 680RD Goat anti-Rabbit IgG (H + L) | LI-COR (926-68071) | 1: 20000 (WB) |
| Normal rabbit IgG | CST (2729) | 1:100 (IP) |

### Mass Spectrometry

For MS sample preparation, cell pellets were treated and collected same as the method in immunoprecipitation. HaloTrap Magnetic Agarose (proteintech, otma) was separated as 25 µL aliquot for each reaction. The beads were washed three times with 500 µL IP lysis buffer, followed by complete removal of supernatant. Then, cell lysates were added to each tubes and incubated at 4 °C for 4 h with gentle rotation. After incubation, beads were intensively washed with IP lysis buffer and eluted in 2 X SDS-loading buffer heated at 95 °C for 5 mins. The samples were submitted to SDS-PAGE and stained with Coomassie brilliant blue solution, and the bands around 250 kDa were cut out, washed three times by $ddH_2O$, destained in 100 µL destaining buffer for 20 mins at 25 °C and this process was repeated once. The gel was then washed by 100% ACN for 15 mins and dried. Then tris(2-carboxyethyl) phosphine (TCEP) was added to reduce protein for 30 mins at 25 °C, followed by addition of iodoacetamide (IAA) solution for 30 mins reaction at 25 °C in dark. The supernatant was discarded and wash by 100% ACN, freeze dried. The sample was then digested by Trypsin solution for 20 h at 37 °C. After centrifugation, supernatant was collected and digested peptides were extracted by the following steps. Extraction solution was added and incubated for 20 mins at 25 °C; this step was repeated once and 100% ACN was added to extracted again. All extracted fractions were combined and freeze dried and desalt by C18 columns. The elutes from columns were freeze dried and injected to MS.

**Mass spectrometry acquisition procedures**

LC-MS/MS data was collected by timsTOF Pro2 mass spectrometer coupled with a nanoElute UPLC system (Bruker Daltonics, Bremen, Germany). The peptides were dissolved in phase A (0.1% formic acid in water). 100 ng of peptides were analysed by C18 column (Aurora Series, 75 μm × 25 cm, Ionopticks, Victoria, Australia) with the gradient set as: 0-48 min, 4-18% solvent B (0.1% formic acid in ACN); 48-55 min, 18-35% B; 55-57 min, 35-95% B; 57-60 min, 95% B with flow rate at 300 nL/min. Peptides were analysed using DDA mode by LC-MS/MS. The parameters set as follows: scan range (m/z) = 300-1500; tims scan range or 1/K0 range (V·s/cm2) = 0.75-1.35; MS1 resolution = 60,000; Target Intensity = 100000; Intensity Threshold=2500; number of PASEF MS/MS scans=6; Total cycle time=1.16s; charge range = 2–5; Isolation Width: 2 m/z (when <800m/z), 3 m/z (when > 800 m/z);

**Mass spectrometry data analysis**

The raw data were processed using PEAKS Studio (version 11, Bioinformatics Solutions Inc., Waterloo, Canada) against O60885 BRD4 in UniProt/SwissProt human (Homo sapiens) protein database. The parameters set as follows: Precursor Mass Error Tolerance: 20.00ppm, Fragment Mass Error Tolerance: 0.05Da, Enzyme: Trypsin, Max Missed Cleavage: 2, Digest Mode: Specific, Peptide Length Range: 5 - 45, Max Variable PTM per Peptide: 3, Fixed Modifications: Carbamidomethylation (+57.02), Variable Modifications: HexNAcylation (ST) (+203.08), Oxidation (M) (+15.99) , Phosphorylation (STY) (+79.97) Database: O60885, Taxonomy: all species, Searched Entries: 1, Deep Learning Boost: Yes, Report Filter: , PSM -10LgP >= 15.0, Proteins -10LgP >= 15.0, Proteins Unique Peptides >= 1.

**Chemical synthesis**

NMR spectra were acquired on Bruker 400 & 500 NMR spectrometer, running at 400 MHz for $^1$H and Bruker 500 NMR spectrometer at 126 MHz for $^{13}$C respectively. $^1$H NMR spectra were recorded at 400 MHz in $CDCl_3$, using residual $CHCl_3$ as the internal standard. $^{13}$C NMR spectra were recorded at 126 MHz in $CDCl_3$ using residual $CHCl_3$ as the internal standard. Thin layer chromatography was performed on Compounds were purified using preparative thin layer flash chromatography (ALUGRAM Xtra, 818333). Mass spectrometry was performed on Agilent LC-MS/MS system consisted of two Agilent 1290 series pumps and auto-sampler, coupled with 6430 triple quadrupole mass spectrometer equipped with and ESI source (Agilent Technologies, Inc., Santa Clara, CA, USA). Unless otherwise noted, analytical grade solvents and commercially available reagents were used without further purification. Unless otherwise noted, chemical starting materials are purchased from Bide pharm without further purification.

**(R)-1-(3-(2-((2-(2-((6-chlorohexyl)oxy)ethoxy)ethyl)amino)-2-oxoethoxy)phenyl)-3-(3,4-dimethoxyphenyl)propyl (S)-1-((S)-2-(3,4,5-trimethoxyphenyl)butanoyl)piperidine-2-carboxylate (OGTAC-1)**

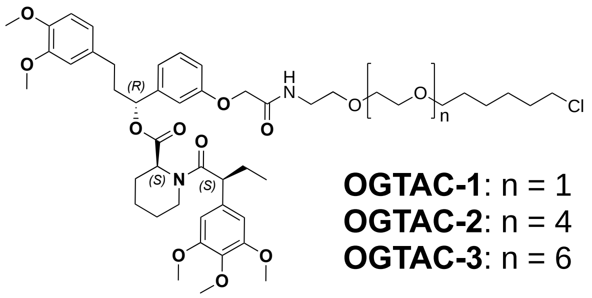


General method: Synthesis of OGTAC-1/2/3 were adapted from literature.[1] Take OGTAC-1 as an example, the 2-[3-[(1R)- 3-(3,4-dimethoxyphenyl)-1-[(2S)-1-[(2S)-2-(3,4,5- trimethoxyphenyl)butanoyl]piperidine-2-carbonyl]oxy-propyl]phenoxy]acetic acid (55.2 mg, 0.079 mmol) (**AP1867**, synthesized according to literature[2,3]) was dissolved in DMF (1.5 mL), 1-[Bis(dimethylamino)methylene]-1H-1,2,3-triazolo[4,5-b]pyridinium 3-oxide hexafluorophosphate (HATU) (35 mg, 0.1 mmol), and DIPEA (50 µL, 0.35 mmol) were added and stirred for 30 minutes. $NH_2$-$PEG_2$-C6-Cl (20 mg, 0.1 mmol) was added. The reaction mixture was stirred overnight. The reaction mixture was extracted with ethyl acetate and water, purified on preparative TLC (ALUGRAM Xtra, 818333), and evaporated under vacuum to give product as clear oil (26 mg, 35 %). Product confirmed by ESI–MS m/z: 899.6 $[M+H]^+$, 921.5 $[M+Na]^+$, 937.6 $[M+K]^+$,; and $^1$H-NMR according to literature.[1] Other probes are synthesised using same reagents just changing $NH_2$-$PEG_2$-C6-Cl to $NH_2$-$PEG_5$-C6-Cl & $NH_2$-$PEG_7$-C6-Cl.

**4-((2-(2-((6-chlorohexyl)oxy)ethoxy)ethyl)carbamoyl)-2-(6-(dimethylamino)-3-(dimethyliminio)-3H-xanthen-9-yl)benzoate (rhodamine ligand)**

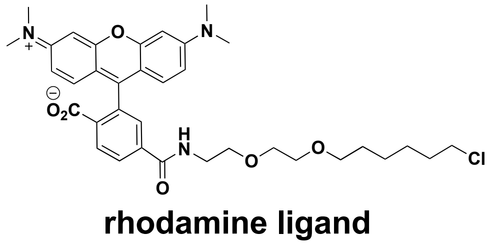

**rhodamine ligand**

This probe was synthesised following the method described in literature.[4] 5(6)-TAMRA NHS ester (14 mg, 0.027 mmol) was dissolved in DMF (2 mL) and ten equivalents diisopropyethylamine (DIPEA) (24 µL) was added to the resultant solution. Then $NH_2$-$PEG_2$-C6-Cl (10 mg, 0.039 mmol) was added to reaction. The reaction was protected from light and reacted for 8 h. Subsequently, the product was dissolved in water and freeze dried, followed by reconstitution in MeOH. The product was purified by PTLC to get final product as dark red powder (15 mg, 86 %). Product confirmed by ESI–MS m/z: 646 $[M+H]^+$.

**(R)-1-(3-(2-((6-((tert-butoxycarbonyl)amino)hexyl)amino)-2-oxoethoxy)phenyl)-3-(3,4-dimethoxyphenyl)propyl (S)-1-((S)-2-(3,4,5-trimethoxyphenyl)butanoyl)piperidine-2-carboxylate (AP1867-C6-NHBoc)**

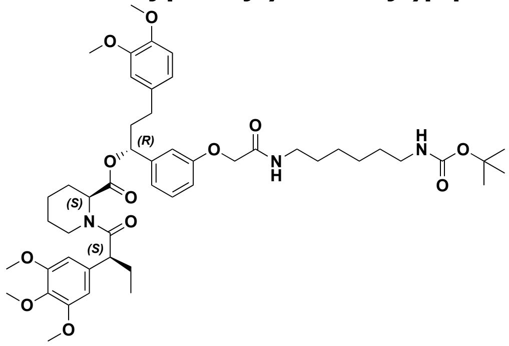


To a round bottom flask (RBF), AP1867 (57.8 mg, 0.08 mmol) was added and dissolved in 2 mL dry DMF. Then DIPEA (70 µL) and HATU (45.6 mg, 0.12 mmol) were added to the solution and stirred for 15 mins. N-Boc-1,6-diaminohexane (25.3 mg, 0.1 mmol) was then added and stirred for another 16 h. The reaction was then partitioned in EtOAc and $H_2O$, extracted with EtOAc and the dried over $Na_2SO_4$. The crude product was purified by PTLC (EtOAc: Hexane= 2 : 1) to have clear oil (33 mg, 51 %). ESI–MS m/z: 691 $[M+H]^+$(without Boc). $^1$H-NMR: δ 7.21 – 7.14 (m, 1H), 6.83 – 6.77 (m, 3H), 6.72– 6.63 (m, 3H), 6.43 (d, J = 2.0 Hz, 2H), 5.67 (dd, J = 8.2, 5.5 Hz, 1H), 5.49 – 5.45 (m, 1H), 4.50 (m, 2H), 3.88 – 3.81 (m, 10H), 3.78 (s, 3H), 3.68 (s, 5H), 3.61 (t, J = 6.7 Hz, 1H), 3.40-3.31 (m, 2H), 3.15-3.06 (td, 2H), 2.86-2.76 (t, 1H), 2.65-2.47 (m, 2H), 2.37-2.28 (m, 1H), 2.15-1.98 (m, 4H), 1.79 – 1.25 (m, 5 H), 1.60 (s, 6H), 1.45 (s, 9H), 0.93 (t, 3H); $^{13}$C-NMR: δ 172.86, 170.73, 168.32, 157.43, 153.30, 148.99, 147.78, 142.47, 136.73, 135.40,133.45, 129.95, 120.31, 119.96, 113.65, 113.13, 111.80, 111.38, 105.07, 75.73, 60.90, 56.43, 56.04, 55.97, 52.20, 50.91, 43.60, 40.70, 39.10,38.36, 31.42, 30.06, 29.83, 29.59, 28.45, 26.92, 26.55, 26.42, 25.44, 21.02, 12.86, 12.69.

**(*R*)-1-(3-(2-((6-(2-((*S*)-4-(4-chlorophenyl)-2,3,9-trimethyl-6*H*-thieno[3,2-*f*][1,2,4]triazolo[4,3-*a*][1,4]diazepin-6-yl)acetamido)hexyl)amino)-2-oxoethoxy)phenyl)-3-(3,4-dimethoxyphenyl)propyl (*S*)-1-((*S*)-2-(3,4,5-trimethoxyphenyl)butanoyl)piperidine-2-carboxylate (OGTAC-4)**

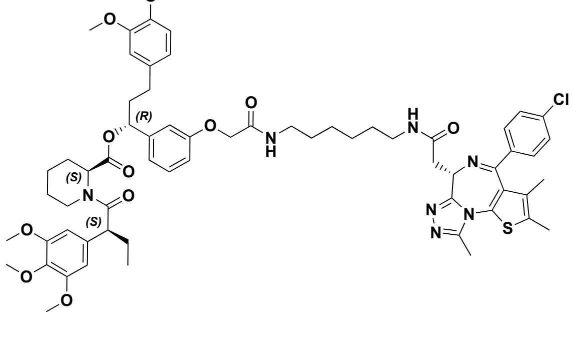


To a RBF, JQ-1 (carboxylic acid, MedChemExpress, HY-78695 ) (13.2 mg, 0.03 mmol) was added and dissolved in 1 mL dry DMF. Then DIPEA (26 µL) and HATU (17 mg) were added to the solution and stirred for 15 mins. **AP1867-C6-NHBoc** (20 mg, 0.03

mmol) was then dissolved in 1 mL dry DMF and added. The reaction was stirred for another 16 h. The reaction was then partitioned in EtOAc and $H_2O$, extracted with EtOAc and the dried over $Na_2SO_4$. The crude product was purified by PTLC (MeOH: DCM= 1 : 30) to have white solid (28 mg, 79 %). ESI–MS m/z: 1175.5 $[M+H]^+$, 1196.5 $[M+Na]^+$. $^1$H-NMR: $^1$H-NMR: δ 7.39 (d, J = 8.4 Hz, 2H), 7.31 (d, J = 8.4 Hz, 2H), 7.21 – 7.14 (m, 1H), 6.83 – 6.77 (m, 3H), 6.72– 6.63 (m, 3H), 6.43 (d, J = 2.0 Hz, 2H), 5.67 (dd, J = 8.2, 5.5 Hz, 1H), 5.49 – 5.45 (m, 1H), 4.63 (t, 1H), 4.50 (m, 2H), 3.88 – 3.81 (m, 10H), 3.78 (s, 3H), 3.68 (s, 5H), 3.61 (t, J = 6.7 Hz, 1H), 3.39-3.30 (m, 2H), 2.86-2.76 (t, 1H), 2.68 (s, 3H), 2.65-2.47 (m, 2H), 2.42 (s, 3H), 2.37-2.28 (m, 1H), 2.15-1.98 (m, 5H), 1.79 – 1.25 (m, 6H), 1.68 (s, 3H), 1.81-1.31 (m, 8H), 0.93 (t, 3H); $^{13}$C-NMR: δ 172.86, 170.71, 168.19, 164.65, 157.44, 153.27, 150.15, 148.95, 147.43, 142.39, 136.68, 135.41,133.47, 132.03, 131.75, 131.30, 130.64, 130.20, 129.90, 128.92, 120.29, 119.86, 113.69, 113.11, 111.79, 111.70, 105.05, 76.9, 70.64, 60.99, 56.39, 56.06, 55.96, 54.41, 52.16, 51.31, 43.57, 38.95, 38.90,38.31, 32.39, 31.38, 29.55, 28.48, 28.45, 26.88, 25.41, 21.00, 13.26, 12.82.
